# Functional recovery from tactile stimulation after perinatal cortical injury is mediated by FGF-2

**DOI:** 10.1101/2022.04.06.487396

**Authors:** Robbin Gibb, Bryan Kolb

## Abstract

The consequences of perinatal brain injury can be devastating but currently there are few effective treatments. We sought to determine if tactile stimulation (TS) or exogenous application of fibroblast growth factor-2 (FGF-2) following injury could reverse the behavioral loss. Infant rats received frontal cortex removals on postnatal day 4 (P4) or a sham surgery. The TS animals received thrice-daily 15-minute bouts of stimulation (Experiment 1) on the day following surgery until weaning. In Experiment 2, treated animals received subcutaneous injections of FGF-2 once daily for one week, postsurgery. Behavioral testing began on postnatal day 60. Brains were later processed for Golgi analysis. We show in Experiment 1, that tactilely stimulating infant rats with perinatal cortical injury stimulates functional recovery and reverses injury-related changes in neuronal morphology in the cerebral cortex. The TS induction of recovery is associated with changes in expression of FGF-2 in both the skin and brain. Direct administration of FGF-2 (Experiment 2) is also effective in facilitating recovery, although not as completely as TS. These results suggest that early behavioral intervention after perinatal cortical injury can stimulate plastic neuronal changes that can underlie functional recovery and that these changes are mediated, in part, through an upregulation of FGF-2.

## INTRODUCTION

Perinatal cortical injury has severe behavioral and anatomical sequelae in both laboratory animals and human infants. For example, rats with cortical lesions on the first days of life have more severe behavioral deficits than animals with similar injuries in adulthood. Furthermore, this poor behavioral outcome is associated with a thin cortical mantle and a general atrophy of dendritic fields in remaining cortical pyramidal cells (Kolb, 1995). Similarly, human children with prenatal and perinatal cerebral injuries are at high risk for chronic behavioral disturbances (Banich et al., 1990: Vargha-Khadem et al., 1985). We therefore asked if there might be environmental treatments that could attenuate the devastating functional consequences of early brain injuries. Because it had been shown that tactile stimulation is effective in stimulating growth in premature infants (Field et al. 1986) and newborn rats (Schanberg and Field, 1987) we decided to evaluate the effect of tactile stimulation on recovery from cortical injury in newborn rats (Experiment 1). This treatment has the advantage of being simple, noninvasive, and it is possible to begin administration soon after the injury.

Fibroblast growth factor-2 (FGF-2, bFGF) is a growth factor that stimulates mitosis and synaptogenesis in brain of both infant and adult animals and promotes survival of neurons following brain damage. A study by Wagner et al. (1999) demonstrated that subcutaneous injection of FGF-2 could stimulate neurogenesis in developing and adult rats. The potential curative property of subcutaneous administration of FGF-2 was investigated in Experiment 2. Interestingly, FGF-2 is also expressed in skin where it is important for proliferation and differentiation of dermal melanocytes, and wound healing. A study by Buntrock et al. (1982) demonstrated that an extract prepared from brain tissue of cattle containing FGF-2 was useful in promoting healing of skin wounds. Knock-out mice lacking FGF-2 protein have been produced and are characterized as having abnormalities in the cytoarchitecture of the cerebral cortex, and delayed wound healing (Ortega et al., 1998). These studies led us to wonder if there might be a connection between the reported benefits of tactile stimulation (TS) and FGF-2 expression in the skin. Increased availability of growth factors produced by skin could conceivably affect brain tissue in a similar manner to subcutaneous injection of exogenous FGF-2. With this focus we chose to decapitate a subset of the TS and non TS animals from Experiment 1 to test for relative expression of FGF-2 and its receptor FGFR_1_ (flg) in frontal cortex and skin.

## SUBJECTS AND TREATMENT PROTOCOLS

### Experiment 1. Tactile Stimulation

The tactile stimulation study was done with 110 rats from 14 litters of animals. Rat pups sustained a frontal lesion or sham surgery on postnatal day four (P4). Half of the litters then received tactile stimulation for two weeks beginning on the day following surgery. This yielded 24 non-TS controls, 28 TS controls, 25 non-TS frontals, and 33 TS frontals. There were approximately equal numbers of males and females in each group. The TS consisted of sequentially stroking individual pups with a soft camel hair paintbrush for 15 minutes 3 X per day. The pups in the TS groups were removed from their mother and placed in a Plexiglas cage that had a 1 cm deep layer of “bed of cobs” on the bottom. The pups were transported to an adjacent room and were given gentle TS with a .5 cm diameter camel’s hair histology brush for 15 min three times daily (9 AM; 1 PM; 4 PM). They were then returned to their mother, having been away from her for no more than 20 min. The TS procedure continued until weaning at postnatal day 21. During the first week of stimulation the animals typically went into REM sleep, as characterized by twitching. By the time the animals reached about 14 days old they had become quite active and the experimenter had to follow the animals with the brush to provide the stimulation. At the time of weaning a subset of TS animals was decapitated and their brains were quickly dissected and frozen for Western blot analysis. A section of skin removed from the rostral portion of the back was also harvested for protein analysis.

### Experiment 2. FGF-2 Treatment

The FGF-2 study was done with 110 rats from 9 litters of animals. Rat pups sustained frontal or sham surgery on P4. Approximately half of the animals in each litter then received subcutaneous injections of 10µg/kg FGF-2 for 7 days beginning the day after surgery. This yielded 10 sham-operated females, 16 P4 females, 16 sham-operated males, and 19 P4 males that were given FGF-2 treatment. There were 8 sham-operated and 18 P4 females; 10 sham-operated and 13 P4 males in the group that received injections of vehicle.

Human recombinant FGF-2 (R & D Systems, Minneapolis MN: # 233-FB) was dissolved in a phosphate buffer solution containing 1mg/ml of bovine serum albumin. The FGF-2 was administered so each rat received 10µg FGF-2/kg body weight. The subcutaneous injections of FGF-2 began the day after surgery and continued for one week. Animals not receiving FGF-2 received placebo injections of the diluent.

## SURGERY

On postnatal day 4 (P4) the pups were removed from the nest and cooled in a Thermatron® cooling chamber until their core temperature reached approximately 20°C. The lesion animals had their scalp opened then the frontal bone carefully removed after it was incised with iris scissors. The medial frontal cortex was then removed bilaterally with gentle aspiration. The tissue targeted for removal was the medial subfield of the prefrontal cortex including Zilles (1985) regions Cg1, Cg3, and PL as well as the medial portion of Fr2 of the motor cortex. After aspiration of the cortical tissue, the animals’ scalp was sutured with silk thread drawn by a very fine needle. The remaining control animals underwent a sham surgical procedure in which the scalp was opened and then sutured closed but the skull was not removed. These animals were identified by removal of the tip of the outer toe on their right rear foot. Beginning on the following day, they were given TS three times daily for the next two weeks or subcutaneous injections of FGF-2 for one week.

## BEHAVIORAL METHODS

### Morris Water Task

Beginning at P60 animals were trained on the Morris Water Task using a similar procedure to that described by Sutherland et al. (1983) based on the original task described by Morris (1981). The maze consisted of a circular pool (1.5 m diameter X 0.5 m deep) with smooth white walls. The pool was filled with approximately 25 °C water mixed with 500 ml of skim milk powder, used to render the water opaque. A clear plexiglas platform (11 × 12 cm) was placed in a constant position inside the pool approximately 30 cm from the pool wall. The water level was adjusted so that the platform was invisible to a viewer outside the pool and to a rat swimming in the water. A trial consisted of placing a rat into the water facing the pool edge at one of four compass locations (north, south, east, or west) around the pool’s perimeter. Within a block of four trials each rat started at the four locations in random sequence, and each rat was tested for four trials a day over five consecutive days. If on a particular trial a rat found the platform, it was permitted to remain on it for 10 seconds. A trial was terminated if the rat failed to find the platform after 90 seconds. Each rat was returned to its holding cage for approximately five minutes before the next trial commenced. The swim path for each rat on every trial was recorded using a Poly Track video tracking system (San Diego Instruments) which tracks the swim path and records the latency, distance and dwell time within each quadrant.

### Whishaw Tray Reaching

Following water task training, animals were trained in a skilled reaching task developed by Whishaw et al. (1991). In this task rats were trained to retrieve chicken feed through metal bars at the front of the Plexiglas training cage (28 cm deep x 20 cm wide x 25 cm high). The front of each cage was constructed with 2 mm bars separated from each other by 1 cm, edge to edge and the floor was constructed of wire mesh. A tray (5 cm deep x 2 cm wide x 1 cm high) containing chicken feed pellets was mounted in the front of each cage. To obtain food, the rats had to extend their forelimbs through the bars, grasp, and retract the food pellet. The food tray was mounted on runners to adjust the distance of the food from the bars. Distance adjustment ensured that each rat could not simply rake the food into the cage. Any pellets that the rat dropped inside the cage were irretrievably lost through the mesh on the floor and the animal would have to reach again. During the first few days the rats were trained in pairs in the reaching cages for a period of one half hour per day. Once reach training commenced, the animals were provided with 15 grams of rat chow daily following the training period. The rats were subsequently trained individually for one half hour per day and then at the end of a two-week training period their performance was videotaped for a five-minute interval. Each time the rat reached through the bars whether or not food was obtained was scored as a “reach” and each time food was successfully returned to the cage and consumed was scored as a “hit”. The percentage of hits to total reaches was then calculated for each animal’s taped performance.

## ANATOMICAL METHODS

### Western Blot

Brain tissue was removed from decapitated animals and placed on ice. Following rapid dissection of frontal cortex, brain samples were placed in microcentrifuge tubes cooled on dry ice. Brain samples were sonicated with 800µl of 1% SDS then aliquoted. All samples were held at –75 ° C until analysis. Samples were diluted 1/20 to determine protein concentration (Bradford Assay) before resolving the protein of interest on 8-12% acrylamide gels (5 µg of protein per well) using SDS-PAGE gel electrophoresis. Frozen skin samples were sonicated with 5mls of 1% SDS. The tissue was then removed and cut into .5 cm squares and resonicated in 5 mls of fresh 1% SDS. After the second sonication the sample was pooled and 1ml aliquots were made and stored frozen at –75C. Gels were blotted on polyvinylidene difluoride (PVDF) membranes. Membranes were blocked for non-specific binding for 1 hr. with 5% non-fat dry milk in Tris-buffered saline and .1% Tween 20. FGF-2 primary antibody (Santa Cruz #sc -7911) was diluted 1:1000 (flg antibody [Santa Cruz #sc-121; 1:1000], in the same solution as was used for blocking. Membranes were incubated in primary antibody for 2 hours followed by 5 washes in PBS (5 mins.) and 1 hour incubation in secondary antibody (HRP-linked donkey anti-rabbit; Amersham #NA934V; 1:5000). Following 5 more PBS washes, FGF-2 protein was revealed with an ECL+ detection kit from Amersham (#RPN2132) and the resulting image was captured on Hyperfilm ECL (Amersham #RPN1647K). Exposed film was imaged with a Kodak digital camera and the blot density was then analyzed using NIH Image software. The membrane was then stained with .1% Commassie Blue (OmniPure EM Science) to reveal all proteins in order to ensure pipetting consistency. A random protein band was selected for each blot and the amount of target protein was calculated as the density of the target / density of random sample protein in the same well. This method allowed compensation for any pipetting errors that may have occurred.

The Western blotting was repeated with a second group of animals in order to replicate the FGF-2 results and to also reveal actin as an additional measure of consistent pipetting consistency. Protein extraction and western blotting was conducted as described by Kovalchuk et al (2018). Membranes were incubated with antibodies against FGF-2 (1:1000; Santa Cruz Biotechnology, Santa Cruz, CA) and actin (1:1000; Abcam, Cambridge, MA). Antibody binding was revealed by incubation with horseradish peroxidase-conjugated secondary antibodies and the ECL Plus TM immunoblotting detection system (GE Healthcare, Piscataway, NJ). Chemiluminescence was detected by the FluoChemHD2 system (ProteinSimple, Santa Clara, CA, USA). Protein loading was visualized by Coomassie brilliant blue R250 staining (BioRad, Mississauga, Ontario, Canada). Images were quantified using Image J Software Package. Statistical analysis was performed using Excel 2013.

### Histological Procedures

At the conclusion of behavioral testing the remaining animals were given an overdose of sodium pentobarbital and intracardially perfused with a solution of 0.9% saline. The trimmed brains were weighed and then immersed whole in 20 mls of Golgi-Cox solution. The brains were then stored (in the dark) in the Golgi-Cox fixative for 14 days before being transferred to a solution of 30% sucrose for seven days. The tissue was cut at 200 µm on a Vibratome™ then developed using a method described by Gibb and Kolb (1998).

### Golgi-Cox Analysis

Layer III pyramidal cells in Zilles’ area Par 1 were traced using a camera lucida at 250X. In order to be included in the data analysis, the dendritic trees of pyramidal cells had to fulfill the following criteria: (a) the cell had to be well impregnated and not obscured with blood vessels, astrocytes, or heavy clusters of dendrites from other cells; the apical and basilar arborizations had to appear to be largely intact and visible in the plane of section. The cells were drawn and analyzed using a Sholl analysis for estimation of dendritic length (Sholl, 1956) was performed. For this analysis a transparent overlay of concentric circles spaced 20 µm apart was placed over the neuron drawing by centering the innermost ring in the middle of the cell body. The number of dendrite-ring intersections was counted for each ring and the total number used to estimate total dendritic length in µm (number of intersections X 20). Five cells were drawn in each hemisphere of each rat. The statistical analyses were done by taking the mean of the measurements on the five cells for each hemisphere of each subject.

Spine density was measured from one apical dendritic branch in the terminal tuft and one basilar terminal branch on five different cells per hemisphere. Spine density measures were made from a segment greater than 10 µm in length, and usually about 50 µm. The dendrite was traced (1000X) using a camera lucida and the exact length of the dendritic segment calculated by placing a thread along the drawing and then measuring the thread length. Spine density was expressed as the number of spines per 10 µm. No attempt was made to correct for spines hidden beneath or above the dendritic segment so the spine density values are likely to underestimate the actual density of the dendritic spines.

## STATISTICAL ANALYSES

All statistical analyses were ANOVA’s performed on Statsview 5® except for Western Blot 2 as noted above. If an ANOVA did not show a significant effect of sex, the data were collapsed across this variable to increase the number of subjects per group and to simplify the analysis.

## BEHAVIORAL RESULTS

### Experiment 1: Morris Water Task

### Latency

Animals with P4 frontal lesions took longer to locate the hidden platform in the water task than did sham operates. Post-operative TS attenuated this impairment. A two-way ANOVA with lesion and treatment as factors showed a main effect of lesion (F(1,66)=88.5, p<0.0001), treatment (F(1,66)=9.17, p=0.0035), and the interaction (F(1,66)=6.34, p=0.014). The stroking treatment had a greater effect on the lesion animals than on the sham-operates giving rise to the interaction (Figure 1).

**Figure 1.**
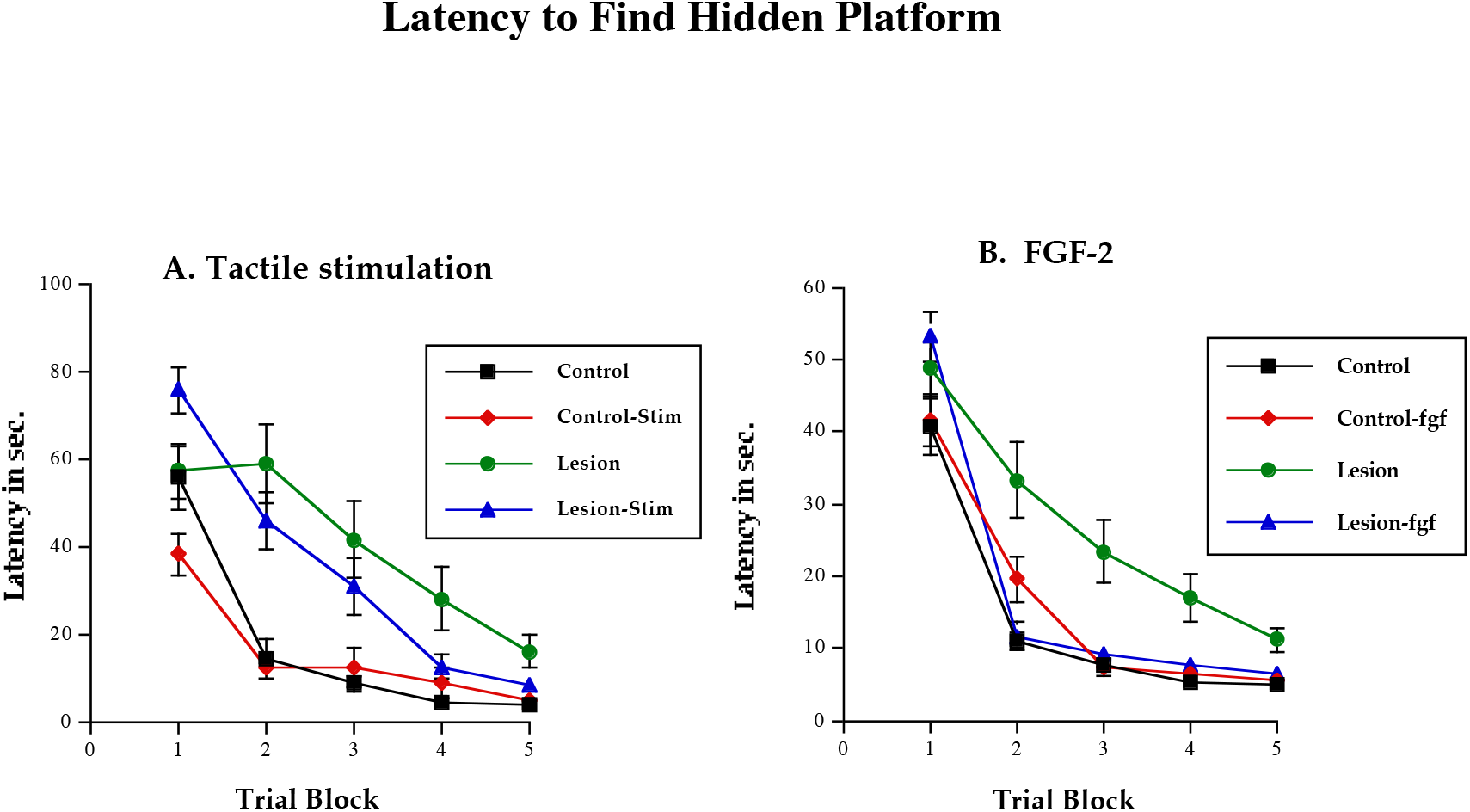
Acquisition curve (latency to find the platform in seconds tested over 5 consecutive days) for Experiments 1 & 2. Data are means±SEs.

#### Distance

Animals with P4 frontal lesions also swam further to find the platform than sham operates. Once again, TS improved the performance of P4 lesion animals in this task (Figure 2). A two-way ANOVA with lesion and treatment as factors showed a main effect of lesion on total swim distance (F(1,42)=9.78, p<.005), and a marginal effect of treatment (F(1,42)=3.8, p<.06). In addition, the Lesion by Treatment interaction was significant (F(1,42)=7.5, p<.01), reflecting the greater effect of the treatment on the performance of the lesion animals.

**Figure 2.**
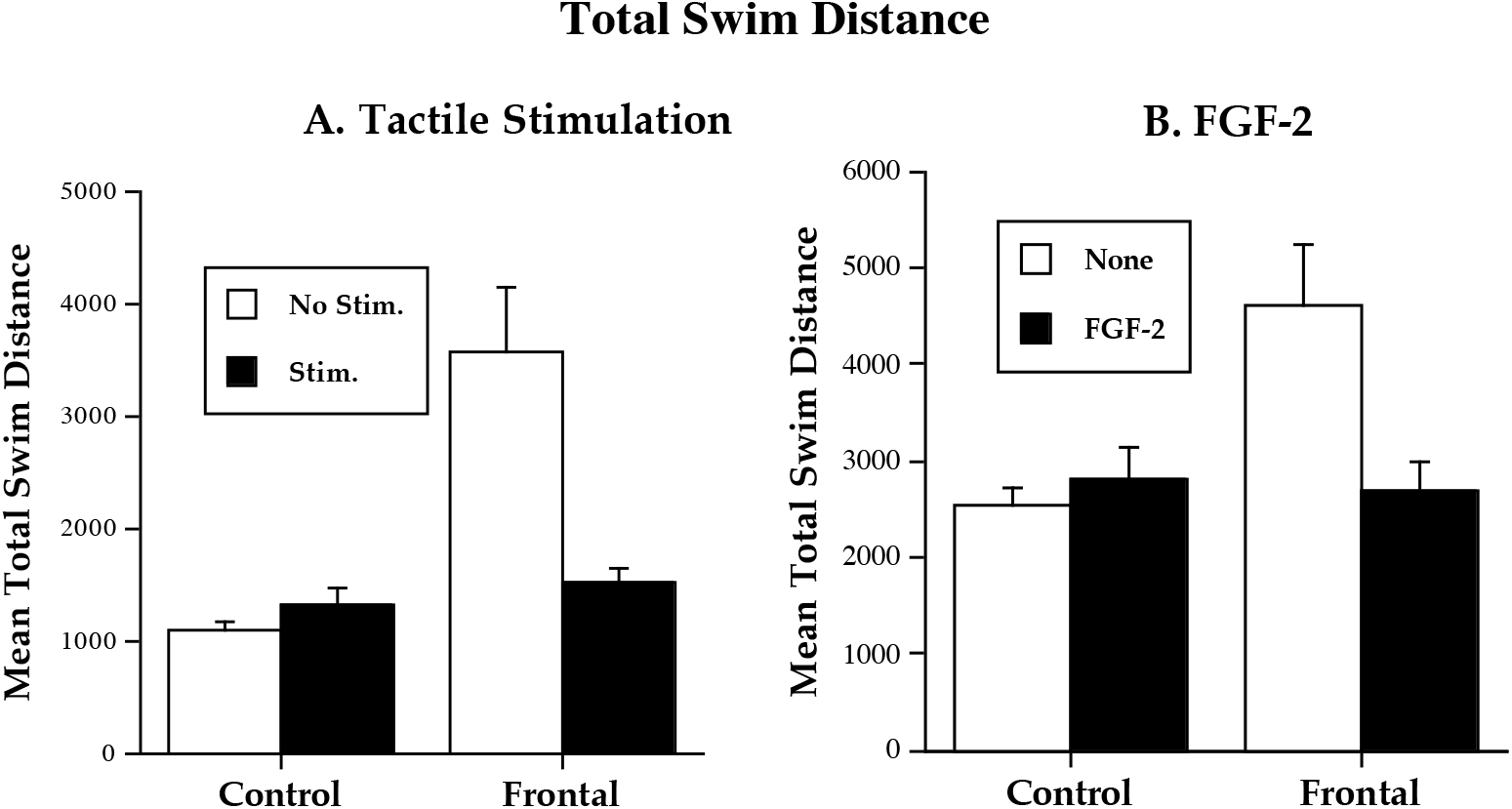
Total swim distance over 5 test days for Experiments 1 & 2. Units are arbitrary computer units. Data are mean±SEs.

### Experiment 2: Morris Water Task

#### Latency

Administration of FGF-2 improved spatial learning in the P4 operates. A two-way ANOVA on escape latency with lesion and treatment as factors showed a marginal effect of lesion (F(1,61)=3.47, p<.07) but no main effect of treatment (F(1,61)= 2.25, p=.14). The Lesion by Treatment interaction was significant (F(1,61)=4.92, p<.05) reflecting the improved performance of lesion animals following FGF-2 administration (Figure 1).

#### Distance

A two-way ANOVA on swim distance with lesion and treatment as factors showed a main effect of lesion (F(1,50)=5.12, p=0.028) but no main effect of treatment (F(1,50)=2.58, p=0.115) and a marginal interaction (F(1,50)=3.52, p=0.066). Once again the interaction reflected the improvement of the lesion animals that received FGF-2 administration (Figure 2).

### Experiment 1: Whishaw Tray Reaching

Lesion animals showed impairments in successfully retrieving pellets for consumption when compared to littermate controls. This impairment was reduced by TS (Figure 3). A three-way ANOVA performed with lesion, treatment, and sex as factors showed a significant main effect of lesion (F(1,68)=89.5, p<.0001), treatment(F(1,68)=8.0, p<.0002), and sex (F(1,68)=5.9, p<.02). None of the interactions were significant (p’s>.48). Posthoc tests showed that both the lesion and control animals benefited significantly from the early experience and that overall, females were better at this test than males (data not shown).

**Figure 3.**
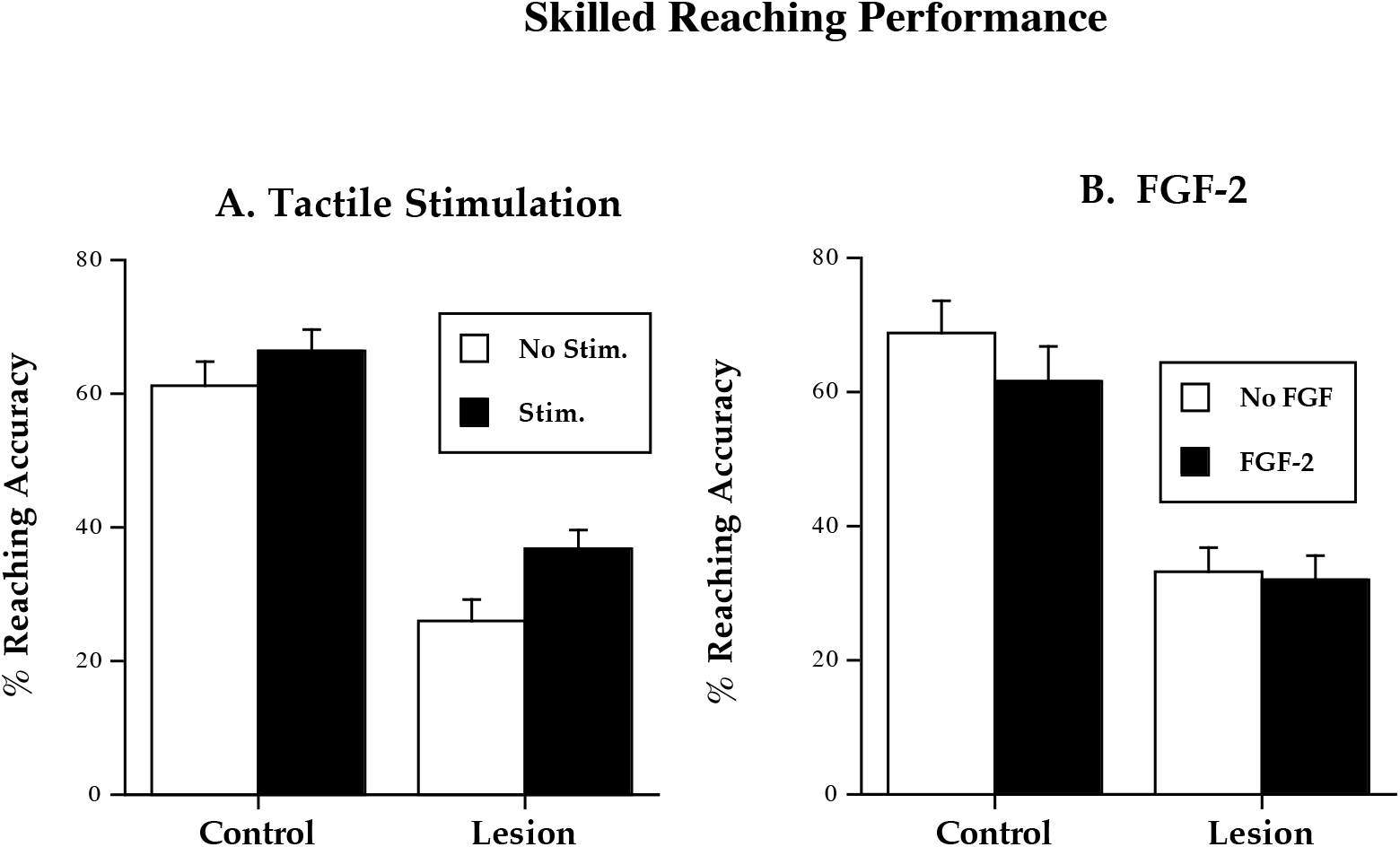
Reaching for Experiments 1 and 2. Bars represent total number of successful reaches /total number of reaches. Data are mean±SEs.

### Experiment 2: Whishaw Tray Reaching

Animals in the lesion group showed deficits in reaching performance compared to the sham-operates and treatment with FGF-2 did not reduce these deficits. A three-way ANOVA with lesion, treatment, and sex as factors showed a main effect for lesion (F(1,95)=61.16, p<0.0001) and sex (F(1,95)=10.77, p=0.0014), but not treatment (F(1,95)=0.297, p=0.59). None of the interactions were significant (p’s>0.21). As with the TS, females performed better on the task then did males across all groups (data not shown).

## ANATOMICAL RESULTS

The frontal lesions were similar in both experiments (Figure 4). The lesions removed the anterior midline cortical tissue including Zilles’ areas Fr2, Cg1, and Cg3. The infralimbic cortex was spared as were the medial and ventral orbital regions. There was no direct damage to the striatum although the anterior striatum was visibly smaller than normal, likely because of the loss of the fibers of passage that would normally course through from the tissue that was removed. The lesions extended posterior to about the level of the septum but did not include the septum or fimbrial regions. For clarity, the anatomical results of Experiments 1 and 2 are described separately.

**Figure 4.**
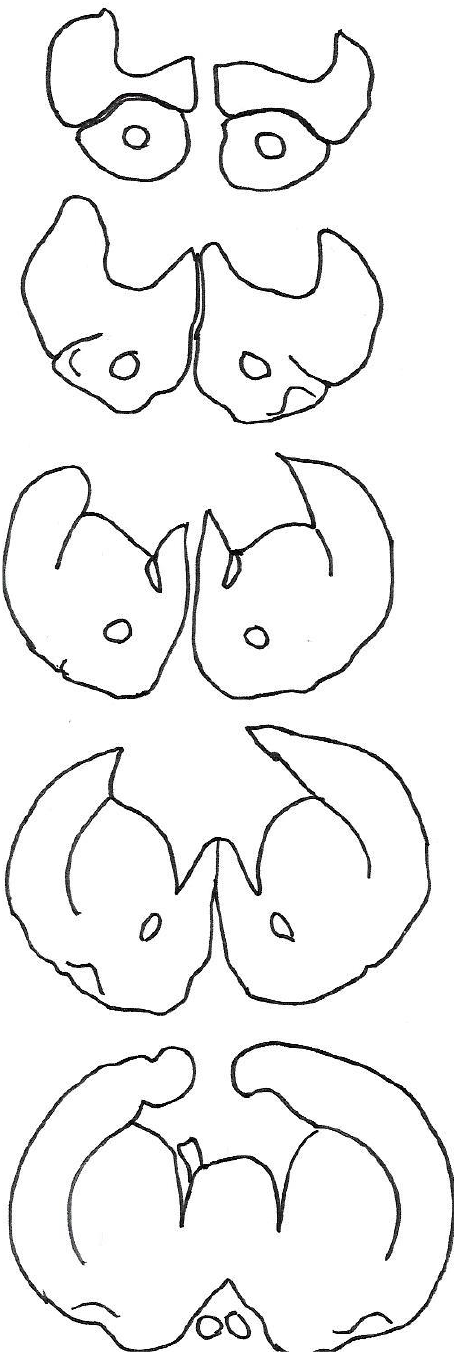
Representative frontal lesion for Experiments 1 and 2.

## EXPERIMENT 1

### Brain weight

TS increased brain weight in both sham and lesion animals (Table 1). A three-way ANOVA on brain weight with lesion, treatment, and sex as factors revealed significant main effects for each factor (F(1,96)=116.2, p<.0001; F(1,96)=3.92, p=.05; F(1,96)=33.9, p<.0001). None of the interactions were significant (p’s>.30)

**Table 1.** Experiment 1: Summary of brain weight

| <u>Group</u> | <u>Brain weight (Males)</u> | <u>Brain weight (Females)</u> |
| --- | --- | --- |
| Sham | 2.055±.03 | 1.883±.04 |
| Sham + stimulation | 2.080±.03 | 1.919±.03 |
| Frontal | 1.750±.03 | 1.637±.03 |
| Frontal + stimulation | 1.812±.04* | 1.699±.02 |

Numbers represent means ± SEs. Brain weight (in gms) shown
\*Differs significantly from same sex frontal group (p<.05)

### Golgi Analysis

#### Dendritic length

Dendritic length was reduced in the apical tree by the lesion and in the basilar tree by the treatment (Table 2). A two-way ANOVA on apical length with lesion and treatment as factors found a significant main effect of lesion (F(1,48)=6.3, p<.02) but not treatment (F(1,48)=0.74, p=0.4). A two-way ANOVA on basilar length found a significant main effect of treatment (F(1,48)=3.99, p=0.05), but not lesion (F(1,48)=0.507, p=0.48) No interactions were significant on either tree (p’s>.4).

**Table 2.** Experiment 1: Summary of dendritic measures

| <u>Group</u> | <u>Apical Length</u> | <u>Basilar Length</u> |
| --- | --- | --- |
| Sham | 1658±34 | 1740±52 |
| Sham + stimulation | 1608±34 | 1632±44* |
| Frontal | 1550±44 | 1700±52 |
| Frontal + stimulation | 1536±32 | 1598±56* |

\*Differs significantly from untreated sham or lesion group, respectively (p<.05).

#### Spine Density

The TS treatment altered spine density in a qualitatively different manner in the sham and lesion animals: it was decreased in the sham operates and increased in the lesion animals. A two-way ANOVA (Lesion X Treatment) on spine density in the apical field of layer III pyramidal cells in parietal cortex revealed main effects of lesion (F(1,184)=37.4, p<.0001), stimulation (F(1,184)=6.5, p<.01), and the interaction (F(1,184)=113.7, p<.0001) (Table 3, Figure 5). Thus, in non-TS animals the lesion reduced spine density whereas in the TS lesion animals there was an increase in spine density relative to control animals.

**Table 3.** Experiment 1: Summary of Group Spine density Apical Spines Basilar Spines

| Group | Apical Spines | Basilar Spines |
| --- | --- | --- |
| Sham | 6.18±0.06 | 6.59±.05 |
| Sham + stimulation | 5.47±0.19* | 5.90±.05* |
| Frontal | 4.92±0.09 | 6.45±.09 |
| Frontal + stimulation | 6.13±0.02* | 6.96±.03* |

Numbers are spines per 10 μm of dendritic length ± SE
\*Differs significantly from untreated sham or lesion group, respectively (p<.05)

**Figure 5.**
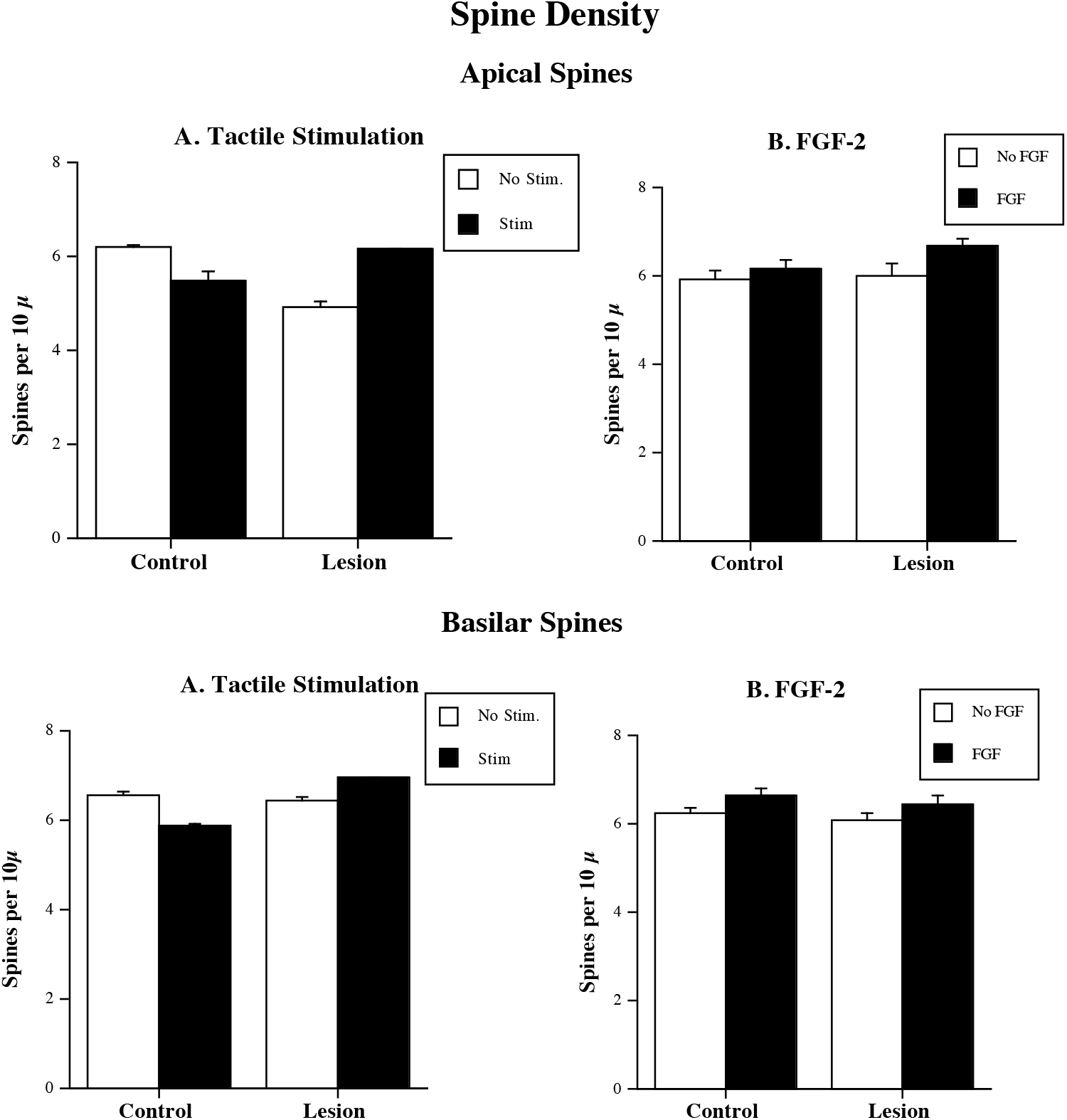
Dendritic spine density measured as average number of spines per 10µm on apical and basilar dendrites in Experiments 1 and 2. Data are mean±SEs.

ANOVA (Lesion X Treatment) on spine density in the basilar field of layer III pyramidal cells in parietal cortex revealed no main effects of lesion or stimulation (p’s>.17), but the interaction of Lesion by Treatment was significant (F(1,184)=37.2, p<.0001), again reflecting the differential effect of the treatment on the lesion animals.

### Western Blot

Frontal cortex samples were harvested from control and lesion animals. In the control animals the sample included all cortical tissue anterior to +3.0 mm ahead of the Bregma. In the lesion animals the sample was taken from approximately the same location but only included the remaining lateral cortex including Frontal area 3 (Fr3) and Agranular insular cortex (AID) according to the nomenclature used by Zilles (1985). The olfactory bulb region was discarded in all samples. Skin samples were taken from just below the neck to mid-torso along the back. Excess hair was quickly removed before the sample was frozen on dry ice. Both brain and skin samples were run to detect the expression of FGF-2 and its receptor FGFR1 (flg) in TS and non-TS animals (Table 4, Figure 6).

**Table 4.**
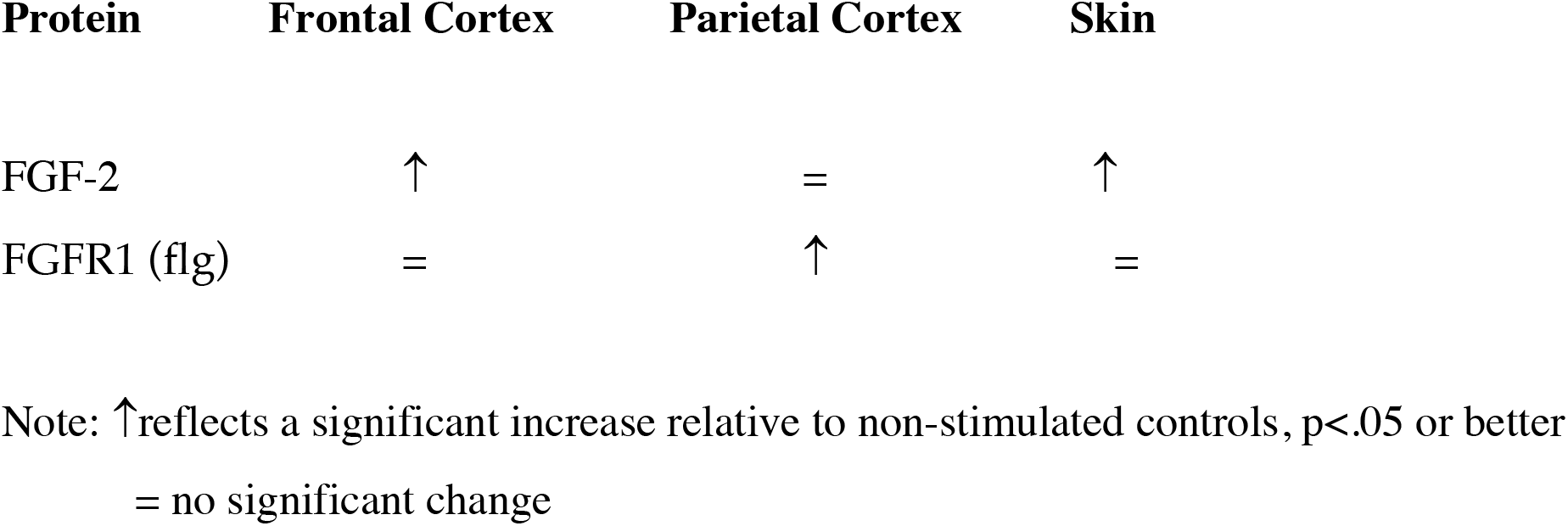
Summary of the findings from the Western Blot experiments.

**Figure 6.**
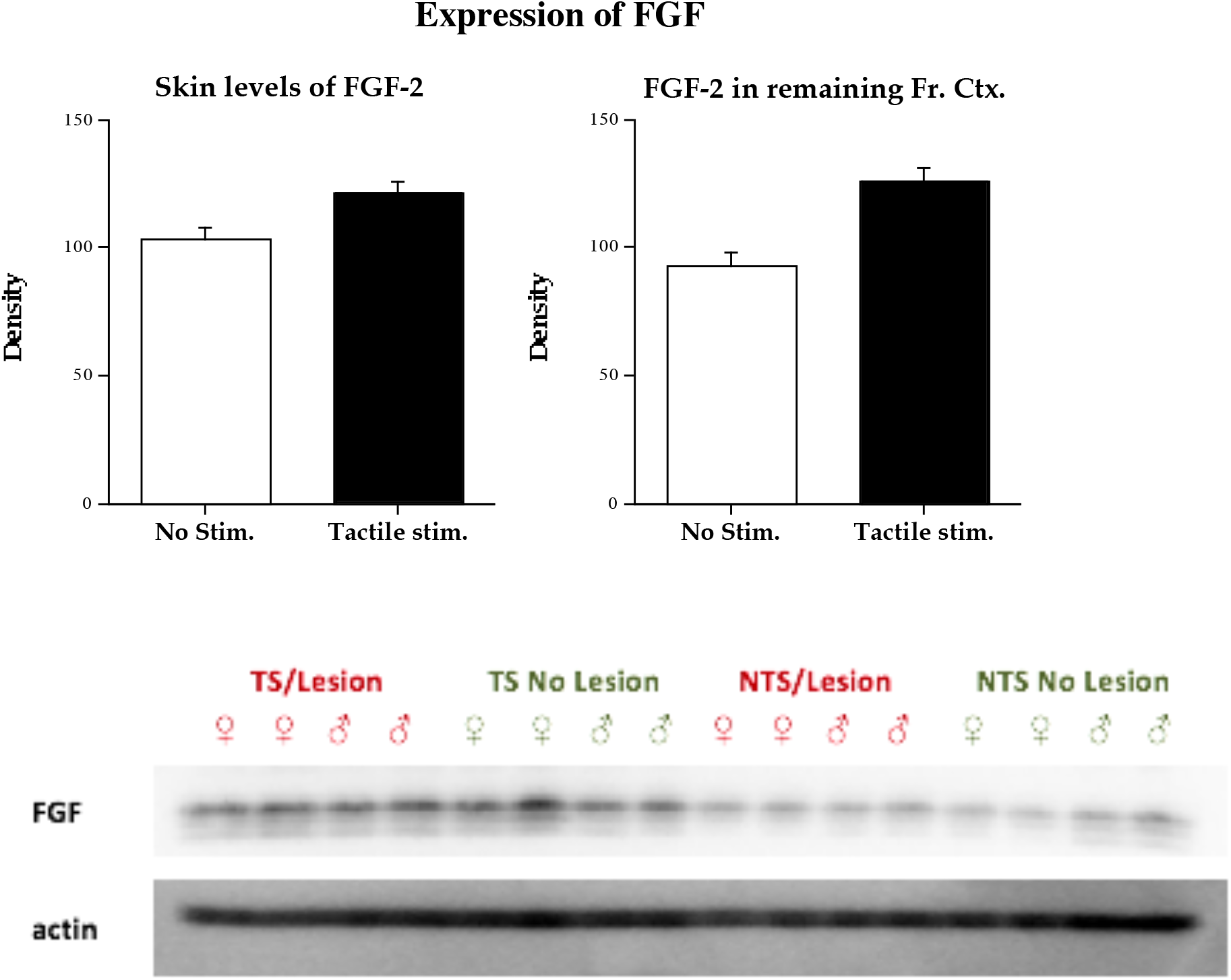
Relative expression of FGF-2 in skin and brain from TS and non-TS animals from Experiment 1. Data are mean±SEs.

An ANOVA looking at the effects of treatment on protein expression of FGF-2 in skin samples of TS and non-TS animals revealed a significant main effect of treatment (F(1,11)=5.37, p=0.041). The TS animals had significantly more FGF-2 in their skin than did the untreated animals. An ANOVA examining flg levels in the skin showed no differences in the levels of this protein in TS or non-TS animals.

An ANOVA on FGF-2 levels in remaining frontal cortex of lesion animals showed a significant main effect of treatment (F(1,6)=6.4, p=0.045). FGF-2 levels were elevated in the TS animals. ANOVA on flg expression showed no differences in TS and non-TS rats (F(1,12)=0.53, p=0.48).

In parietal cortex there were no differences observed in FGF-2 expression. ANOVA showed no effect of treatment (F(1,12)=0.024, p=0.88). There was an increase in expression of the flg protein, however (F(1,12)=4.99, p=0.047).

## EXPERIMENT 2

### Brain Weight

In contrast to the effects of TS, there was no significant effect of FGF-2 administration on brain weight (Table 5). A three-way ANOVA on brain weight with lesion, sex, and treatment as factors revealed significant main effects for lesion and sex but not treatment (F(1,89)=110.2, p<.0001; F(1,89)=12.8, p=.0006; F(1,89)=.132, p=.72). The Lesion X Sex X Treatment interaction was significant (F(1,89)=5.3, p=.02) and reflected a decrease in brain weight in the female control group following FGF administration.

**Table 5.** Experiment 2: Summary of brain weight

| <u>Group</u> | <u>Brain weight (Males)</u> | <u>Brain weight (Females)</u> |
| --- | --- | --- |
| Sham | 1.929±.02 | 1.829±.03 |
| Sham + FGF-2 | 1.972±.02 | 1.809±.02 |
| Frontal | 1.709±.05 | 1.596±.03 |
| Frontal + FGF-2 | 1.704±.03 | 1.643±.03 |

Numbers represent means ± SEs. Brain weight (in gms) shown.

### Golgi Analysis

#### Dendritic length

The P4 lesion reduced dendritic length in the apical tree as did the FGF-2. ANOVA on apical length with lesion and treatment as factors showed a significant effect of lesion (F(1,80)=12.7, p=0.0006) and treatment (F(1,80)=15.9, p=0.0002) but not the interaction (Table 6). Similar effects were seen in the basilar tree. A two-way ANOVA revealed a significant main effect of lesion (F(1,80)=8.74, p=0.04) and treatment (F(1,80)=32.8, p<0.0001) and a trend in the interaction (F(1,80)=3.62, p=0.06). The interaction reflected a reduction in dendritic length following P4 lesion in the untreated animals as compared to the increase in dendritic length observed in the FGF-2 treated animals.

**Table 6.**
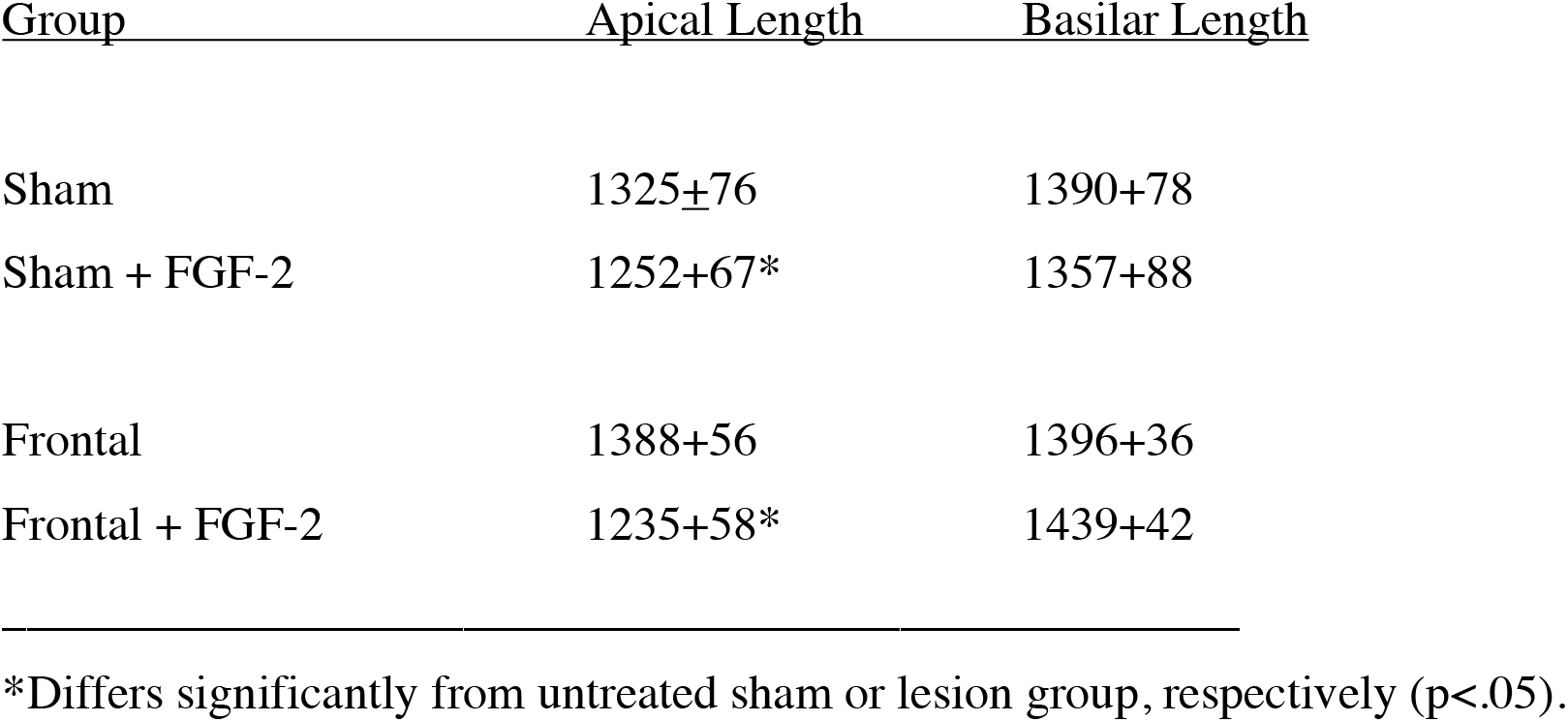
Experiment 2: Summary of dendritic measures

| <u>Group</u> | <u>Apical Length</u> | <u>Basilar Length</u> |
| --- | --- | --- |
| Sham | 1325±76 | 1390±78 |
| Sham + FGF-2 | 1252±67* | 1357±88 |
| Frontal | 1388±56 | 1396±36 |
| Frontal + FGF-2 | 1235±58* | 1439±42 |

\*Differs significantly from untreated sham or lesion group, respectively ( $p < .05$ ).

#### Spine Density

Treatment with FGF increased spine density in the apical spines (Table 7, Figure 5). A two-way ANOVA (lesion, treatment) on the apical tree showed no main effect for lesion (F(1,80)=1.81., p=0.18), nor the interaction (F(1,80)=1.16, p=0.28) but there was a main effect of treatment (F(1,80)=4.69, p=0.033). Posthoc tests (Fisher’s PLSD, p<.05) showed a significant treatment effect in the lesion but not the control animals. In the basilar tree ANOVA revealed no main effect of group (F(1,88)=0.15, p=0.70)n or the interaction treatment (F(1,80)=0.01, p=0.92, but there was a main effect of treatment (F(1,80)=4.62, p=0.035). Posthoc tests showed that neither control nor lesion group differed significantly from their respective FGF-treated groups (p>.10).

**Table 7.** Experiment 2: Summary of Spine density

| Group | Apical Spines | Basilar Spines |
| --- | --- | --- |
| Sham | 5.95±0.20 | 6.24±.15 |
| Sham + FGF-2 | 6.17±0.20 | 6.65±.15 |
| Frontal | 6.00±0.29 | 6.10±.16 |
| Frontal + FGF-2 | 6.68±0.16* | 6.47±.18 |

Number of spines per 10 μm dendritic length ± SE
\*Differs significantly from untreated frontal group.

## DISCUSSION

There were three novel findings of these studies. First, both the TS and FGF-2 significantly reduced the behavioral impairments after cortical injury. Injury to the frontal region produced a marked deficit in behavioral performance on both tasks relative to sham-control animals and this deficit was significantly reduced by the TS and by administration of FGF-2. In fact, the brain-injured rats showed such an improvement in performance that they were able to perform almost as well as control animals in the spatial navigation task. Skilled reaching was improved by TS but not by FGF-2. This latter result may have been a problem of effective dose of FGF-2. In a recent study (Waite and Kolb, 2003) FGF-2 (10µg/kg) solution was made fresh daily and a significant behavioral recovery was observed on skilled reaching. In the current study the FGF-2 was made fresh at the beginning of the treatment and then stored at 4° C. We must note, however, that although the TS was beneficial, the lesion animals still showed significant impairments.

Second, tactile stimulation altered brain morphology and did so differently in sham and cortically injured animals. The overall findings were that: 1) tactile stimulation reduced dendritic length in the basilar field; and 2) frontal cortical lesions significantly reduced dendritic length in the apical field. The TS-related decline in dendritic length was unexpected but the lesion-related decline replicates our previous findings (Kolb and Gibb, 1990). FGF-2 administration also decreased dendritic length in both the apical and basilar fields. The frontal lesion also caused a reduction but in the basilar tree the reduction was less dramatic in the FGF-2 treated animals than in the untreated operates.

The analysis of dendritic spines led to an unexpected result: Sham-operated animals that received tactile stimulation had a significant decline in spine density in both the apical and basilar fields whereas frontal-operates with similar stimulation had a significant increase in spine density. Thus, a similar experience differentially affects the intact and injured brain. The reduction in spine density with early experience has precedents (Kolb et al., 2003) but the opposite effect of experience on the injured brain is a novel finding. A similar finding was noted in the FGF-2 experiment. Although FGF-2 did not affect spine density in the apical tree, in the basilar field there was trend for FGF-2 treatment to reduce spine density in controls and increase spine density in the frontal cortex operates.

Third, tactile stimulation increased the expression of FGF-2 in both the skin and brain. We hypothesized that the change in cortical morphology was related to an increase in one or more neurotrophic factors and in particular to FGF-2. FGF-2 is known to pass the blood-brain barrier and is made not only in the brain but also in the skin in response to injury or stimulation. We therefore used Western blotting to investigate the effects of tactile stimulation on FGF-2 expression in skin and cortex, and found an increase in both places in the stimulated groups. The presence of an increase in FGF-2 expression in both skin and cortex of tactilely-stimulated rats, combined with the functional improvement and associated anatomical changes, suggests that the behavioral treatment acted at least in part, through its actions on FGF-2 expression. The fact that direct administration of FGF-2 was also able to enhance functional outcome, although somewhat less effectively, is consistent with this hypothesis. It is well established that behavioral treatments can influence the production of neurotrophic factors (Carro et al., 2000; Berchtold et al., 2002; Vaynman et al., 2003). What has not previously been recognized, however, is that the same behavioral treatment can differentially affect the intact and injured brain. It is not immediately obvious why a neurotrophic factor treatment would have differential effects on the normal and injured brain but the injured brain presumably has altered regulation of many factors, including gene expression that could be affected by neurotrophic factors.

There is a rich behavioral literature showing that early experiences in infancy can permanently affect the brain and behavior of adult rats (Denenberg and Whimbey, 1963; Denenberg et al., 1967; Levine, 1967; Levine et al., 1967, Meaney et al., 1987; Meaney et al. 1988). The current study suggests that early intervention after cerebral injury is especially powerful in influencing brain and behavioral development. The importance of early intervention after cortical injury cannot be underestimated. We have shown elsewhere that complex-environment versus cage housing can stimulate recovery after early frontal lesions but only if it begins at weaning (Kolb et al., 2003). In fact, even four months of complex rearing has a minimal effect upon recovery from early frontal lesions if it does not begin until adulthood. Furthermore, the magnitude of the effects observed in the current study after just two weeks of tactile stimulation are greater than those we have found after four months of complex rearing. We should note, however, that the addition of FGF-2 to complex housing experience has greater effects on functional recovery after cortical injury in adulthood (B. Kolb and A. Witt-La Jeunesse, unpublished observations). Perhaps the failure of behavioral therapies later in life to facilitate functional recovery is that they are not effective in sufficiently raising FGF-2 levels. Or, it seems equally likely that the tactile stimulation affected more than just FGF-2 and whatever other factors that are altered by the tactile stimulation may also be necessary to facilitate functional recovery.

Finally, we would be remiss if we did not address the question of whether our tactile stimulation treatment is related to its effects on stress. There is a large literature showing that exposure to stressful experiences, such as brief, repeated removal from the mother during development, can affect cognitive processes in adulthood and senescence (Meaney et al., 1987; 1988). We have shown, however, that such procedures have no beneficial effect on functional recovery from early injury and, in addition, may actually retard recovery (Gibb and Kolb, 2005). Furthermore, such procedures also do not reverse the injury-related drop in spine density (Gibb and Kolb, 2005). The tactile-stimulation procedure in the current study may actually have reduced stress insofar as we noted that during the first week or so of tactile stimulation the animals typically entered into a sleep pattern that was punctuated by twitching, suggesting that the animals were in rapid eye movement sleep. It thus seems plausible to hypothesize that whereas brief periods of neonatal stress may be beneficial in the adult animal, it may not be advantageous to the animal suffering from perinatal brain injury.

## Acknowledgements

This research was funded by grants from NSERC of Canada to BK and graduate awards from NSERC and AHFMR to RG. The authors thank Grazyna Gorny for her assistance in anatomical analyses, Tara Moroz, Mike Oorth, and Connie Greschner for their assistance in the stimulation study and Anna Kovalchuk, Kurt Schlachter, Lesley Schimanski, and Brian West for their help in the FGF-2 study.

